# A mechanical origin for implantation defects in embryos from aged females

**DOI:** 10.1101/2025.09.29.679218

**Authors:** Kate E. Cavanaugh, MJ Franco-Oñate, Diana J. Laird, Patrick W. Oakes, Ricard Alert, Orion D. Weiner

## Abstract

Women over 35 experience a marked reduction in fertility. The origin of these fertility defects appears to reside in the implantation capacity of the embryo itself, but the mechanistic basis of this impairment is not well-understood. Here, we identify a core mechanical defect in embryos from aged mothers that impairs the process of implantation. Using mouse models, we find that reproductive aging yields increased contractility in the extra-embryonic trophectoderm, the outer epithelial tissue responsible for mediating uterine attachment and embryo implantation. This hypercontractile state elevates tissue surface tension and viscosity in the blastocyst, culminating in defective spreading during implantation. Enhanced contractility is necessary and sufficient for this age-related defect in implantation, and early embryo mechanics can be used to predict successful implantation for embryos from both young and aged mothers. Our work represents a potential foundation for improving embryo selection in Assisted Reproductive Technologies to resolve age-related defects in female fertility.

## Introduction

After the age of 35, women have only a 5–10% chance of achieving a full-term pregnancy, and more than 60% of miscarriages occur around the time of embryo implantation^1–4^. Age-related infertility was originally attributed to the uterine environment, in which the endometrium was thought to become increasingly stiff, fibrotic, and less receptive to implantation^5,6^. However, clinical outcomes from In Vitro Fertilization (IVF) challenge this view, as pregnancy rates in older women are largely rescued when using donor embryos from younger women^7^. These IVF data suggest that the embryo and not the uterus is the dominant driver of age-associated infertility. Age-related changes in embryo mechanics may compromise an embryo’s ability to undergo the complex morphogenetic events required for successful uterine integration^8–13^, but the underlying embryonic defects remain poorly understood. Because the average age of maternal childbirth continues to rise worldwide, uncovering how maternal aging alters the earliest stages of embryo development represents a pressing and unresolved challenge with significant implications for both basic biology and the improvement of IVF outcomes.

A major barrier in the IVF field lies in selecting blastocysts that will successfully lead to a full-term pregnancy^14–17^. Despite decades of refinement, current screening approaches yield only about a 40% success rate for younger women and 15% for women over the age of 35^18,19^. The gold-standard assessment of embryo quality remains the gross morphology of the blastocyst structure^20,21^. Here, IVF grading typically evaluates how effectively the primitive placental Trophectoderm epithelial layer encloses the fluid-filled cavity surrounding the compact, fetal Inner Cell Mass (ICM) (**Fig. 1a**). However, morphological grading is subjective, context-dependent, and does not always predict embryo viability. More recently, additional screening methods have included the biopsy of Trophectoderm cells for aneuploidy testing^22,23^. Although chromosomal abnormalities are prevalent in human embryos, many embryos labeled “euploid” still exhibit implantation defects^23^. This highlights that chromosomal status, while important, does not fully capture the developmental competence of an embryo. Together, these limitations underscore a critical gap: neither morphology nor genetic screening reliably captures the functional viability of an embryo.

**Fig. 1.**
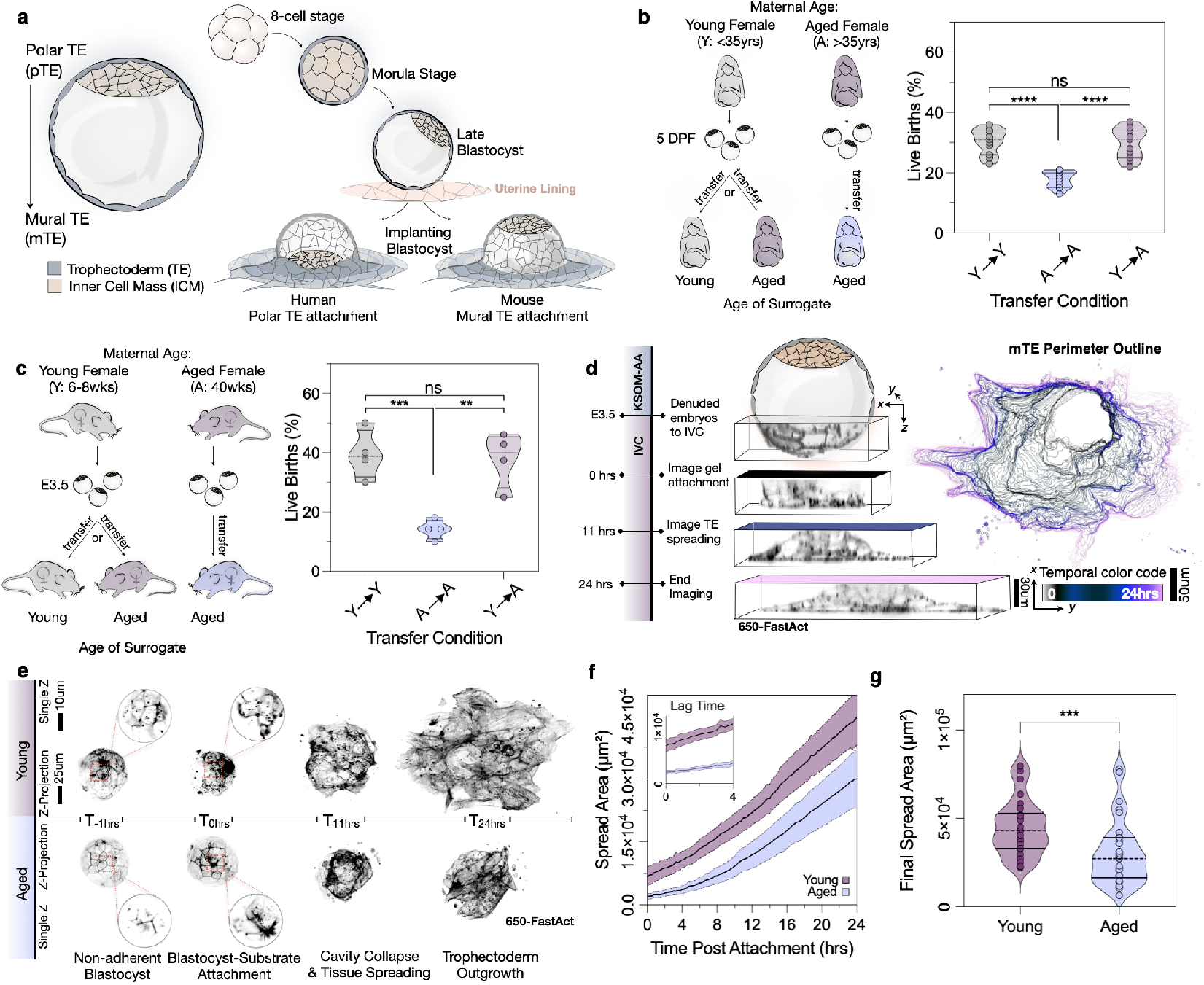
Embryos from aged mothers display low implantationx. a) Schematic of blastocyst structure with the Inner Cell Mass (ICM) (tan) sitting within the blastocyst and surrounded by the Trophectoderm tissue (gray). The ICM sits at the Polar Trophectoderm (pTE) region, with the opposite side being the Mural Trophectoderm (mTE) (left). Schematic of distinct developmental checkpoints spanning the 8-cell, morula, blastocyst, and implanting blastocyst stage. Species-specific differences occur at the site of implantation: the mouse embryo implants at the mural Trophectoderm, whereas the human embryo implants at the polar Trophectoderm (right). b) Schematic of Uterine Transfer workflow in Assisted Reproductive Technologies parsed from the HFEA^24^. Embryos are transferred individually to the uterus. Embryos from Young or Aged females are transferred to either Young or Aged surrogate mothers (Y->Y n = 318,975 transferred embryos; A->A n = 34,255 transferred embryos; Y->A n = 482,730 transferred embryos). Violin plots show a reduction in age-related infertility that is rescued with embryos from younger women following transfer to older surrogates. p values from Mann-Whitney test. c) Schematic of NSET workflow to simulate Assisted Reproductive Technologies. Embryos from Young or Aged females are transferred to either Young or Aged surrogate mothers (N=4 for all transfer conditions, Y->Y n = 38 transferred embryos; A->A n = 35 transferred embryos; Y->A n = 36 transferred embryos). Violin plots show a reduction in age-related infertility that is rescued with developmentally younger embryos when transferred to older females. Two tailed p values from N-1 Chi-Square test. d) Diagram showing the experimental workflow for In Vitro Implantation Assays. E3.5 Blastocysts are denuded and transferred from KSOM-AA to IVC media that contained 650-FastAct to visualize the process of implantation on collagen-coated polyacrylamide gels. 24 hours post-implantation, the embryos are fixed (left). Views of the blastocyst in Z show mural Trophectoderm attachment, blastocyst cavity collapse, and spreading over the 24 hours post-implantation (middle). Mural Trophectoderm perimeter outlines show the evolution of mTE spreading over 24 hours. Perimeter outlines are color coded based on time post-implantation (right). Scale bars are 30μm and 50μm, as indicated. e) Representative images of Young and Aged implanting blastocysts over the 24 hours post-implantation. Blastocysts are initially non-adherent, after which they extend protrusions to mediate attachment, cavity collapse, and tissue spreading. Inlays display a single z-plane to visualize mural Trophectoderm protrusions at the substrate surface. Images of implanting blastocysts are displayed as a z-projection. Scale bars are 10 μm and 25 μm, as indicated. f) Graph of the evolution of the Trophectoderm’s spread area over 24 hours of implantation. Inlay shows a zoom of the first 4 hours of implantation, displaying an age-specific delay in attachment and spreading (Young N = 4, n = 26 implanting blastocysts; Aged N = 4, n = 31 implanting blastocysts). Lines and shading show mean and 95% Confidence Interval. g) Violin plot depicted total spread area post-implantation for both maternal conditions (Young N = 4, n = 26 implanting blastocysts; Aged N = 4, n = 31 implanting blastocysts). Maternally aged embryos show significant reductions in final spread area. p values from Mann-Whitney test.

Here, we investigate the developmental origins of age-related infertility by combining approaches from classical mammalian embryology, developmental cell biology, and biophysics. By applying engineering and biophysical approaches to reproductive health, our work reveals a mechanical origin of age-related implantation defects: hyperactive contractility in mouse embryos from older females impairs multiple developmental events, including uterine implantation. Mouse models replicate age-related defects seen in humans, as assayed via pregnancy outcomes and *in vitro* implantation assays. Here, aging of the mother slows Trophectoderm spreading on synthetic gel substrates. Heightened embryo contractility is both necessary and sufficient for the age-related implantation defects. This peri-implantation defect can be explained by increased surface tension and viscosity of the embryo, which oppose spreading during implantation similarly to how they oppose the wetting of a droplet on a 2D surface. We trace these mechanical changes back to compaction at the 8-cell stage, a key morphogenetic checkpoint during which developmental tempo was accelerated in maternally aged embryos. Together, these data reveal a mechanical signature of early mouse embryos that predicts implantation outcomes for embryos of both young and aged females, providing a potential criterion for embryo grading to improve IVF outcomes.

## Results

### Mouse models replicate human age-associated implantation defects

A decline in human age-associated fertility can be observed in public datasets of IVF blastocyst-uterine transfers (**Fig. 1b**)^24^. This assisted reproductive technology sought to improve fertility outcomes through the collection of eggs from gonadotropin-stimulated donor females following fertilization and transfer to the uterus of a surrogate mother. When donor embryos were collected from younger women (less than 35 years of age) and transferred to similarly young surrogates, a 32% live birth rate was achieved in this IVF procedure. When embryos were collected from older females (beyond 35 years of age) and transferred to older surrogates, the live birth rate dropped to 18%, recapitulating the sharp decline in female fertility as women’s reproductive organs rapidly age after 35 years. Older surrogates receiving embryos from younger female donors exhibited a rescue of fertility to 31% live birth rate, indicating that an older woman’s uterus is receptive to embryo implantation. These data suggest that the origin underlying maternal age-associated infertility is the egg or early embryo, but the basis of this defect is not known.

We leveraged the mouse as a model to investigate the embryonic origin of age-related infertility with a focus on implantation, the stage at which most pregnancy failures occur in humans. The early mouse embryo is amenable to *ex utero* culture, concurrent long-term quantitative microscopy, genetic manipulation by microinjection or electroporation, live-imaging using cell-permeable dyes to visualize protein localization, and mRNA-encoded fluorescent protein expression^25–30^. Murine lineage specification and early preimplantation morphogenetic checkpoints are also analogous to human embryogenesis^31^. Furthermore, the mouse embryo undergoes implantation through similar mechanisms to humans, albeit with notable differences in post-implantation morphogenesis (**Fig. 1a**)^32–35^.

To test the effects of maternal age on pregnancy outcome in mice while modeling the clinical practice used in humans, we leveraged the well-established technique of Non-Surgical Embryo Transfer (NSET)^25,36^. Non-invasive insertion of experimental blastocysts into the uterine horn of a pseudopregnant female by NSET enabled subsequent implantation and full-term pregnancy, after which litter sizes were tracked. This scheme enabled us to quantify live birth rates similar to published IVF datasets (**Fig. 1b**). In this experimental workflow, we used young sexually mature male mice (<35 weeks of age) to remove the variable of paternal age. We bred males to females from two distinct age groups: young (6-8 weeks old) and aged (40+ weeks old), roughly equivalent to women in their 20’s and 40’s, respectively^37,38^ (**Fig. 1c**). Embryos pooled from younger female donors (which we termed as “young embryos”) were cultured to blastocyst stage and transferred to young surrogate females, which resulted in a 41% live birth rate. Embryos from older female donors (which we termed as “aged embryos”) transferred to aged surrogates, resulted in an 18% success rate, recapitulating the age-related decline in fertility seen in humans. Like their human counterparts, aged mice were rescued to a fertility rate of 42% when they were recipients of young embryos, consistent with an embryonic origin of age-related infertility in both mice and humans. We subsequently focused on a more experimentally tractable system to probe the effect of maternal age on embryo implantation.

The biophysical mechanisms underlying implantation mediated by the trophectoderm remain largely unexplored due to technical limitations of visualizing *in utero* processes. We overcame this barrier by combining biomechanical measurements with previously-established and highly efficient *in vitro* implantation assays^39,40^. This *ex utero* platform harnesses synthetic collagen-coated polyacrylamide (PAA) gels and specialized IVC media to mimic the physiochemical nature of the receptive maternal tissue during embryo-substrate attachment and spreading. Extending *ex vivo* implantation to mouse embryos, we achieved over 90% attachment rates of embryos to the gel substrate, similar to previous reports^39,40^.

To visualize spreading dynamics during trophectoderm invasion in our *in vitro* assays, we performed live confocal microscopy of E4.5 blastocysts stained with 650-FastAct for 24 hours post-implantation^25^. To limit phototoxicity over this extended period, we limited our imaging to the future implantation site at the mural Trophectoderm (mTE) region of the blastocyst to capture its attachment and outgrowth 24-hours post-adhesion (**Fig. 1d, Supp. Movie 1**). We defined the initiation of implantation (t_0_=0hrs) as the transition between a coherent trophectoderm epithelia and the appearance of protrusive trophectoderm cells at the basal substrate (**Fig. 1e in-lays**). By imaging actin dynamics every 15 minutes over 24 hours, we captured the dynamics of implantation, including initial blastocyst-substrate attachment, cavity collapse, and full trophectoderm outgrowth (**Figs. 1d, 1e, Supp. Movie 1**). Embryos from young mothers displayed efficient spreading dynamics, characterized by a relatively rapid transition between attachment, spreading, and outgrowth (**Fig. 1f**). While maternally aged embryos initially attached to the substrate, their spreading resulted in a significant reduction in the final trophectoderm contact area at 24 hours post attachment (**Fig. 1g**). These data indicate a dampening of implantation potential that arises with advanced maternal age.

### Enhanced contractility is necessary and sufficient for maternal age-related implantation defects

What is the origin of reduced trophectoderm implantation in embryos from aged mothers? Cellular contractility is known to be a primary regulator of tissue spreading in other contexts, including the invasion of cancer aggregates^41,42^. Mouse embryos exert contractile pulling forces at cell-substrate adhesions to mediate outward spreading of the trophectoderm during implantation^32^. Here, contractility acts to increase cell-substrate interactions to promote spreading. However, contractility can also act in cell-cell adhesion to limit tissue spreading. As such, it is not a priori clear whether increasing contractility should promote or hinder tissue spreading. We therefore probed whether alterations in contractility underlie the defective spreading in embryos from aged females. To investigate whether blastocyst contractility plays a functional role in embryo implantation, we used pharmacological regulators of actomyosin-based contractility (**Figs. 2a, 2b**).

**Fig. 2.**
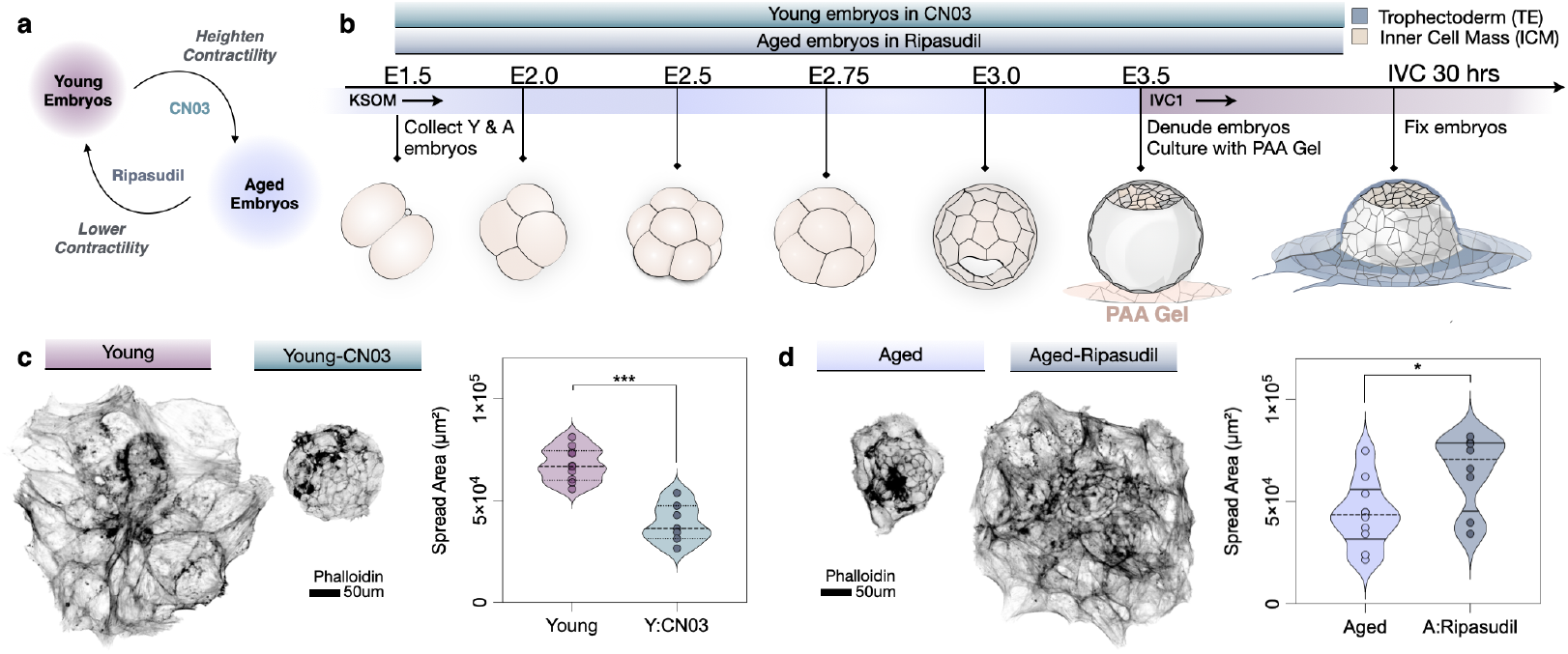
Contractility is necessary and sufficient for maternal age-related implantation defects. a) Schematic of contractile inhibitor pathways to regulate potential trophectoderm spreading behaviors. We hypothesized that RhoA activator CN03 would phenocopy aged embryos, while ROCK inhibitor Ripasudil would rescue aged spreading behaviors. b) Schematic of experimental workflow and the culture conditions for testing the necessity and sufficiency of contractility for maternal age-related implantation defects. Embryos are collected at E1.5, after which Young embryos are cultured in KSOM-AA with or without CN03 (Rho-based contractility activator), and Aged embryos are cultured in KSOM-AA with or without Ripasudil (Rho Kinase-based contractility inhibitor). Embryos are then denuded and transferred to IVC media for In Vitro Implantation assays and fixed 30 hours post-implantation. c) Representative images of Young (left) and Young-CN03 treated (right) embryos fixed and stained for actin. Violin plot shows a reduction of spread area in Young-CN03 treated embryos (Young N = 9, n = 43 implanting blastocysts; Young -CN03 N = 5, n = 25 implanting blastocysts). Scale bar is 50μm. p values from Mann-Whitney test. d) Representative images of Aged (left) and Aged-Ripasudil treated (right) embryos fixed and stained for actin. Violin plot shows a rescue of spread area in Aged-Ripasudil treated embryos (Aged N = 4, n = 27 implanting blastocysts; Aged-Ripasudil N = 3, n = 16 implanting blastocysts). Scale bar is 50μm. p values from Mann-Whitney test.

We first harnessed the cell-permeable RhoA-activator CN03 (**Figs. 2a, 2b**). RhoA is a master regulator of contractility that functions by activating actin polymerization and myosin-mediated contractile force generation. CN03 potentiates RhoA activity by inhibiting GAP-associated GTP hydrolysis to increase the level of GTP-bound RhoA within 2-4 hours^43^. We added CN03 to embryo culture medium starting at the 2-cell stage and then assessed the effects on implantation potential. Gel-implanted embryos were fixed at 30hrs post-implantation to ensure adhesion of experimental blastocysts and proper spreading as observed in our live-imaging (**Figs. 1e, 1g**). Young embryos treated with ectopic RhoA activator CN03 displayed a striking reduction in spread area analogous to maternally aged embryos (**Fig. 2c**).

Enhanced contractility was sufficient to impair trophectoderm spreading in embryos from younger females, mimicking the defects observed in embryos from older mothers. To probe whether enhanced contractility is required for the maternal age-related defect in implantation, we attempted to rescue trophectoderm spreading by reducing contractility in aged embryos (**Figs. 2a, 2b**). We cultured maternally aged embryos in the FDA-approved Y-compound derivative, Ripasudil, which modulates myosin-mediated contractility through inhibiting the activity of Rho-Kinase, ROCK^44,45^. Ripasudil-treated maternally aged embryos exhibited rescued spread area in our *in vitro* implantation assays, indicating a necessary role for actomyosin contractility in the spreading defects in aged embryos (**Fig. 2d**).

We finally confirmed blastocyst contractility as a function of advanced maternal age by immunostaining E4.5 blastocysts with the tension sensor, VD7, which recognizes cell-cell junctions under strain^46^ (**Supp. Fig. 1a**). The mural Trophectoderm region in the aged condition exhibited significant increases in normalized junctional VD7, consistent with increased tissue tension in the site of future implantation. Maternally aged embryos also exhibited increased normalized junctional Non-Muscle Myosin isoform IIb (MYH10) and phosphorylated Myosin Light Chain (pMLC) (**Supp. Figs. 1b, 1c**), both indicators of increased tissue tension.

Because increased contractility in embryos from older females was necessary and sufficient for the observed spreading defects, this indicates a causative role for contractility in dampening embryo implantation mechanics. Our data suggest an optimal regime of contractility for cell spreading. While a certain level of contractility is necessary for outward tissue migration, further increases in contractility (through aging or pharmacological manipulation) hinder embryonic implantation.

### Advanced maternal age impairs blastocyst wetting

We next turned to physical models of tissue spreading to better understand how enhanced blastocyst contractility produces age-related implantation defects. Based on previous work^33,41,48^, we describe implantation using active wetting theory, which models the 3D blastocyst structure as an active liquid droplet spreading on a 2D surface, such as the uterine lining (**Fig. 3a**). Active wetting theory predicts that tissue spreading depends on the competition between blastocyst surface tension and exerted cell-substrate forces. These two forces can have opposite effects; surface tension of the blastocyst acts to minimize contact with the substrate, while cell-substrate forces promote blastocyst spreading (**Fig. 3b**).

**Fig. 3.**
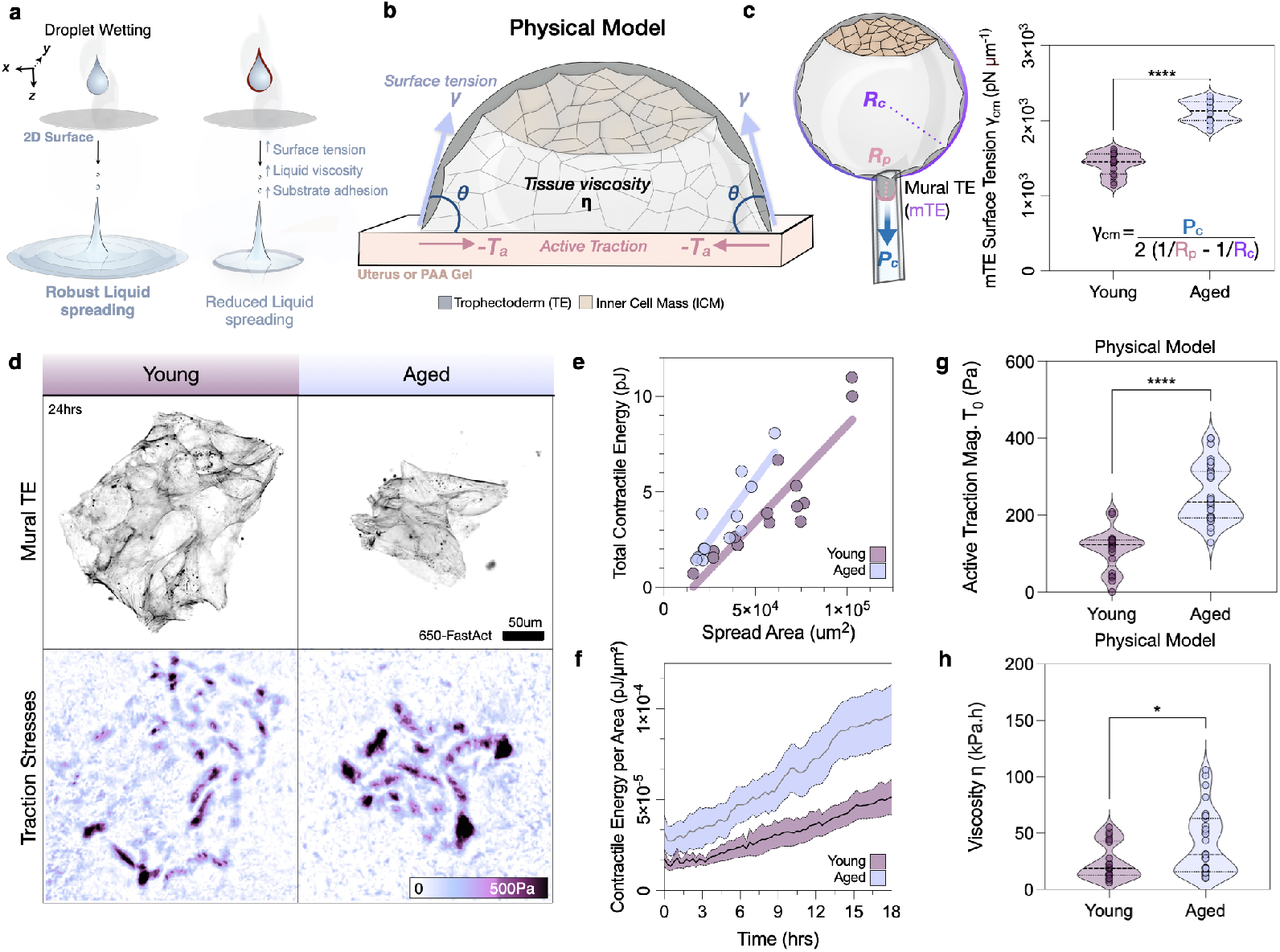
Advanced maternal age dampens blastocyst wetting. a) Droplet spreading is a function of a liquid’s surface tension, adhesion between the liquid and the substrate, and viscosity within the liquid. These parameters collectively determine a droplet’s wetting behavior. When extrapolated to the blastocyst, key parameters dictating a tissue’s spreading behavior include surface tension, cell-cell adhesion, and cell-substrate adhesion. b) Schematic of an implanting blastocyst as a droplet with viscosity *η* whose spreading is driven by active traction forces ***T***_a_ and opposed by surface tension *γ*. The inward-pointing traction forces on the substrate (-***T***_a_) correspond to regions of cell pulling. c) Schematic of Micropipette Aspiration Assay for measuring surface tension of E4.5 Blastocysts (above). The glass pipette aspirates a portion of the mural Trophectoderm region. Violin plot of average surface tension measurements of E4.5 between the two maternal conditions (Young N = 3, n = 15 blastocysts; Aged N = 3, n = 15 blastocysts). p values from Mann-Whitney test. d) Representative images of implanted trophectoderm stained for 650-FastAct (black) along with calculated traction stresses (purple) for both maternal conditions. Scale bar is 50 μm. e) Plot with each dot representing a single embryo, analyzing Total Contractile Energy (pJ) as a function of the trophectoderm’s spread area (um^2^) with simple linear regression to show trends. Aged embryos show higher Contractile Energy per Area compared to younger maternal controls (Young N = 3, n = 18 implanted embryos, R^2^ = 0.81; Aged N = 3, n = 13 implanted embryos, R^2^ = 0.77). f) Contractile energy per Area as calculated by Traction Force Microscopy between both maternal conditions (Young n = 18 implanted embryos; Aged n = 13 implanted embryos). Maternally aged embryos show higher strain energy per area values across the 18 hours post-implantation. Bold lines represent the mean and are bounded by shaded SEM. g) Plot showing Maximal Traction *T*_0_ in both maternal conditions, as determined by fitting the physical model to the data, with higher maximal traction in the aged condition (Young n = 21 implanted embryos; Aged n = 31 implanted embryos). p values from Mann-Whitney test. h) Viscosity is higher in embryos from aged mothers, as determined by fitting the physical model to the data (Young n = 21 implanted embryos; Aged n = 31 implanted embryos). p values from Mann-Whitney test.

We hypothesized that the defective spreading of blastocysts in the aged condition could arise from changes in mechanical forces pre-implantation. To test this hypothesis, we performed direct mechanical measurements of late E4.5 blastocysts via Micropipette Aspiration Assays. As expected, maternally aged blastocysts exhibited increased surface tension at the mural trophectoderm (**Fig. 3c, Supp. Fig. 2a, Supp. Movie 2**). This increased surface tension indicates elevated mTE tissue tension, which could restrict tissue spreading during implantation.

In addition to elevated surface tension, an increase in cell contractility in embryos from aged females could also lead to stronger cell-substrate applied traction forces^41^. To measure cell-substrate forces during *in vitro* implantation^32^, we embedded fluorescent beads in our synthetic substrate PAA gels (**Supp. Fig. 2b**) and applied the biophysical technique of Traction Force Microscopy (TFM). Bead trajectories indicate the substrate deformation and can be used to calculate the applied traction stresses that result from cellular forces at cell-substrate adhesions^41,49,50^. TFM provides a powerful approach to determine the tissue’s contractile state during implantation^32^.

We observed a surprising age-related hyperactivation of contractility within the migrating trophectoderm front in aged embryos (**Fig. 3d, Supp. Movie 3**). Analysis of traction profiles across multiple samples revealed strong fluctuations in active traction forces. This observation was consistent with a recent study that reported intermittent tractions applied by mouse embryos on extracellular matrix^32^. For embryos of the same spread area, aged embryos showed an increase in total contractile work performed, consistent with hypercontractility in embryos of advanced maternal age (**Fig. 3e**). Maternally-aged implanted embryos exhibited a heightened contractile energy per unit spread area over 24 hours (**Fig. 3f**).

Our data indicate that the hypercontractility of embryos from aged mothers leads to an increase in both surface tension and cell-substrate tractions. To assess their impact on embryo spreading, we used our active wetting model (**Fig. 3b**), which is based on force balance, ***∇*** · *σ* + T_a_ = 0, between cell-substrate active tractions T_a_ and internal forces in the tissue, captured by its stress tensor *σ* (**Supplementary Note**). As in our measurements (**Fig. 3d**), our model assumes that tractions are maximal at the tissue leading edge and decay inwards with a decay length *L*_c_ *≈* 15 *μ*m (see Methods) (**Supp. Fig. 2c**). In the model, internal stresses are due to tissue flow *v* with viscosity *η*: *σ*_*αβ*_ = *η*(*∂*_*α*_*v*_*β*_ + *∂*_*β*_*v*_*α*_), where Greek indices denote spatial components. Moreover, following wetting theory, we account for the embryo’s surface tension *γ* through a Young-Dupré condition at the tissue’s leading edge: *σ*_*rr*_(*R*) = −*γ* cos *θ*, where *r* denotes the radial component, *R* the spread radius, and *θ* the contact angle^48^ (**Fig. 3b, Supp. Fig. 2d**).

We then solved the model and fitted it to the area spreading dynamics of all embryos (**Supplementary Note, Supp. Fig. 2e**). The results reveal that, in addition to higher surface tension (**Fig. 3c**) and higher tractions (**Fig. 3g**), aged embryos also exhibit a higher tissue viscosity (**Fig. 3h**), which contributes to slower spreading dynamics to hinder implantation. These analyses support the role of age-related alterations in tissue mechanics in driving defective embryo implantation.

To further validate our modeling results, we assessed the mechanical properties of the tissue using cell shape as a proxy for tissue viscosity^51,52^. Previous work established the dimensionless shape index *q*, defined as the ratio of the cell perimeter over the square root of its area, as an indicator of the mechanical state of a tissue^53,54^ (**Supp. Fig. 2f**). For non-motile cells, values of *q* below the threshold *q*_c_ = 3.8 indicate tissues behaving like a solid, with largely hexagonal cell shapes. Higher values of q indicate a fluid state of the tissue. Cells exhibiting higher shape indices exhibit more elongated and irregular shapes, which yields a lower tissue viscosity. To apply this approach, we analyzed cell shape indices of the mural Trophectoderm region in all embryonic conditions. In all cases, we found *q >* 3.8 (**Supp. Figs. 2f, 2g**), consistent with the tissue being fluid, as assumed in our model. However, in embryos from younger mothers, shape indices were on average higher, indicative of a lower viscosity that may facilitate spreading. In contrast, in aged embryos, shape indices were lower, suggesting a heightened tissue viscosity consistent with the fitted values from our wetting model. This increase in tissue surface tension and viscosity appear to arise from hyperactivation of contractility, as CN03-treated young embryos phenocopied the aged condition (**Supp. Figs. 2g, 1h**).

Together these data indicate that maternally aged embryos exhibited heightened actomyosin contractility that increases regional mTE surface tensions at the future implantation site. Higher cellular contractility leads to increases of mTE cell-medium surface tension (**Fig. 3c**), mTE cell-substrate tractions (**Figs. 3d, 3g**), and tissue viscosity (**Fig. 3h**). These data support an implantation defect arising from impaired blastocyst wetting, in which a heightened tissue viscosity hinders spreading.

### Compaction metrics predict implantation potential in embryos from both young and aged females

Contractility acts as a key driver in multiple morphogenetic stages preceding implantation^8,27,28,55,56^. For example, at the 8-cell compaction stage, compaction proceeds as endogenous contractility flattens the embryo’s outer surface^57^. Often considered the first major morphogenetic checkpoint in the embryo, the timing and extent of compaction is dependent on RhoA-mediated contractility, which is reflected in an embryo’s morphology^30,57^. Embryonic compaction is critical for cells to integrate mechanical stimuli and initiate placental programming in the future trophectoderm lineage, which carries out implantation.^58–60^ We sought to leverage morphogenesis as a metric to create a non-invasive classifier to capture an embryo’s mechanical potential in implantation. The logic behind this experimental framework was: (1) mechanically competent embryos are more likely to accomplish full implantation, and (2) mechanical properties can be identified during Compaction at the 8-cell stage using non-invasive methods.

To perform morphokinetic analyses of preimplantation development, mouse embryos from young and aged females were cultured *ex utero* starting from the 2-cell stage onwards, similar to previous work^61^. We specifically examined developmental tempo, or the dwell time within each stage from the 4-cell, 8-cell, and to the 16-to 32-cell stage when the embryo cavitates for blastocyst formation (**Fig. 4a, Supp. Movie 4**). We define the initiation of each developmental stage as the resolution of cell division from the previous cell cycle, with the stage’s resolution being the initiation of the next subsequent cell division. This approach enables us to assay the developmental timing of maternally aged embryos and document alterations in morphogenesis that may underlie age-associated infertility.

**Fig. 4.**
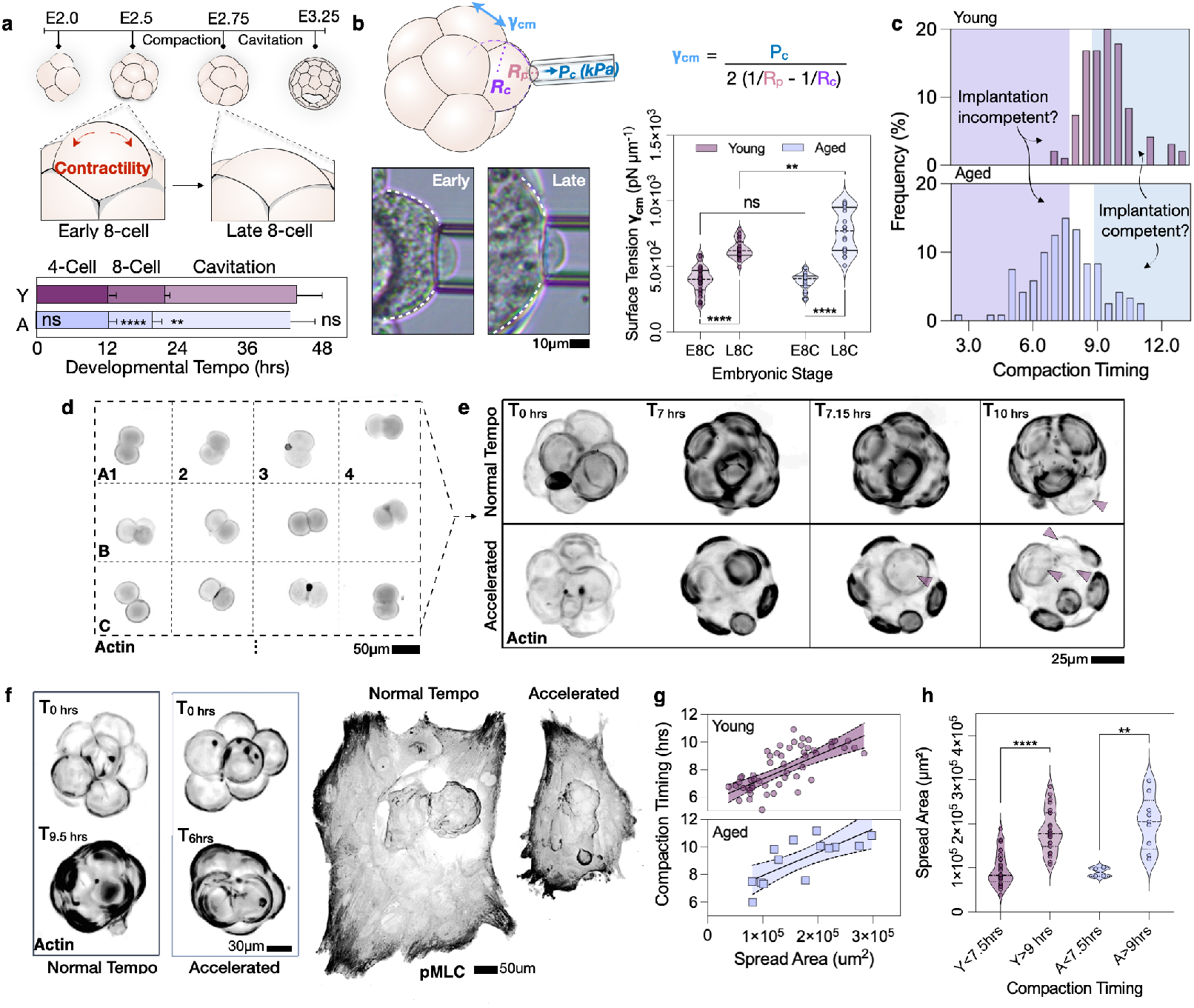
Compaction metrics predict implantation potential. a) Schematic of developmental timing from the 4-cell stage through early blastocyst cavitation. At the 8-cell stage, contractility drives compaction and flattening of the embryo’s outer surface. Below are the quantifications of developmental tempo for the 4-cell, 8-cell, and cavitation stages. Maternally aged embryos display a significant reduction in the timing of the 8-cell stage (N=4, Young n = 95 embryos; Aged n = 119 embryos). p values from Mann-Whitney test. b) Schematic and representative images of Micropipette Aspiration Assays between early and late 8-cell stages, where the pipette aspirates the exposed apical region of the cell to calculate cortical surface tension. Analyses of cellular surface tensions show maternal age-associated increase in tension at the late 8-cell stage (Young N = 8, n = 44 Early 8-cell embryos, 26 Late 8-cell embryos; Aged N = 5, n = 24 Early 8-cell embryos, 19 Late 8-cell embryos). p values from Mann-Whitney test. Scale bar is 10μm. c) Histogram of compaction tempo between both maternal conditions (N=4, Young n = 95 embryos; Aged n = 119 embryos). Boxes highlight the expected implantation incompetent and implantation-competent windows with respect to compaction timing. d) Schematic of the indexing workflow for correlating compaction metrics with implantation potential. Embryos are stained for 650-FastAct and indexed at the 2-cell stage, imaged over the period of compaction, and then selected based on their developmental tempo to undergo In Vitro Implantation. Scale bar is 50μm. e) Representative images from timelapse imaging of embryos stained for 650-FastAct showing Normal (top) and Accelerated (bottom) compaction timing. Embryos in the Normal condition show significantly longer times to the first cell division into the 16-cell stage. Purple arrows denote cell divisions. Scale bar is 25μm. f) Representative images of embryos live-imaged at the 8-cell stages (black, 650-FastAct) undergoing compaction (left) and their representative fixed, implanted embryos (black, pMLC staining) (right). Scale bars are 30μm and 50μm, as indicated. g) Plot of compaction timing as a function of final Trophectoderm spread area 48 hours post-implantation for both maternal conditions (N=3, Young n = 56 implanted blastocysts; N=2, Aged n = 14 implanted blastocysts). Bold line represents simple linear regression and are bounded by 95% Confidence Interval (Young R^2^ = 0.40; Aged R^2^ = 0.55). h) Violin plots from (f) show spread area binned by their compaction timing of either >9hrs or <7.5hrs. Both Young and aged conditions show significant reductions in spread area if their compaction timing falls within the <7.5hrs compaction category (N = 3: Young n = 23 >9hrs, 29 <7.5hrs; N = 2: Aged n = 9 >9hrs, 5 <7.5hrs).

Time-lapse imaging revealed a significant abbreviation of the 8-cell stage in embryos from aged females. In embryos from younger females, the dwell time during compaction stages was 9.5 hours on average, consistent with previous work assessing developmental tempo during the 8-cell stage^57^. In contrast, maternally aged embryos accelerated through the 8-cell stage an average of two hours faster than younger control embryos (**Fig. 4a, Supp. Fig. 3a**). While we also find an age-dependent lengthening of the period to cavitation, we find no significant differences in total developmental timing between young and aged maternal conditions (**Supp. Fig. 3a**).

We confirmed contractility at the 8-cell stage using the direct mechanical measurement of cortical tension via micropipette aspiration (**Fig. 4b, Supp. Movie 5**)^57,62^. As predicted by their accelerated progression through the 8-cell stage, maternally aged embryos showed significantly increased cortical tensions compared to young control embryos post-compaction (**Fig. 4b**). These data reveal an accelerated developmental tempo in embryos from advanced maternal age, leading to precocious compaction driven by aberrant cytoskeletal dynamics and heightened cortical tension in 8-cell stage embryos.

We hypothesized that this mechanical signature at compaction stages would be indicative of later trophectoderm function and outcomes during implantation. In this experimental scheme, we first harness the natural heterogeneity within embryos’ compaction timing to generate a classifier that is predictive of implantation outcomes *in vitro* (**Fig. 4c**).

To link compaction metrics of each embryo to its implantation outcome, we modified our *in vitro* implantation assay by culturing embryos on silicone grids so that each embryo could be indexed (**Fig. 4d**). Cultured embryos were imaged at 5-minute intervals from the 4-to the 16-cell stage, after which we calculated compaction metrics before processing indexed embryos for downstream *in vitro* implantation assays. We segregated embryos for implantation assays by two temporal conditions: 1) compaction timing >9 hours, similar to the younger control embryos, and 2) compaction timing <7.5 hours, similar to the maternally aged condition (**Fig. 4e, Supp. Movie 6**). Within these two binned conditions, we also measured the blastomere contact angles as a metric for each embryo’s contractile state and endogenous cellular force balance (**Supp. Fig. 3b**).

Two interconnected mechanical parameters at the 8-cell stage were predictive of implantation potential in our extended pre- and peri-implantation culture. We observed that the trophectoderm outgrowth spread area at late blastocyst stage correlated with the earlier developmental tempo over the period of compaction (**Fig. 4f**) as well as with the average inter blastomere contact angles just before cell division into the 16-cell stage (**Supp. Fig. 3c**). Embryos that exhibited accelerated compaction and more acute cell-cell contact angles were statistically more likely to restrict spreading of their trophectoderm, resulting in reduced implantation potential *in vitro*. These correlations held for embryos from both young and aged mothers, enabling us to harness natural compaction heterogeneity to generate a classifier that is predictive of outcomes during *in vitro* implantation (**Figs. 4g, 4h, Supp. Fig. 3d**). While earlier mechanical differences linked to implantation defects become more common at advanced maternal age, our results highlight the value of leveraging developmental tempo to identify implantation-competent embryos.

## Discussion

In this study, we uncovered a core mechanical defect in embryos from aged females that underlies their impaired implantation competence. We leveraged this knowledge to identify the small subset of embryos from older mothers that are competent for implantation. Using mouse models, we found that maternal reproductive aging triggers an increase in contractility in the extra-embryonic trophectoderm, decreasing the ability of these embryos to implant both *in vivo* and *in vitro* (**Figs. 1, 3**). Enhanced contractility is both necessary and sufficient to drive the age-related defects of implantation failure (**Fig. 2**). Contractility acts to increase effective tissue viscosity, impairing implantation of embryos from older females (**Fig. 3**). Furthermore, non-invasive assays for altered mechanics (timing of embryo compaction, blastomere contact angles, direct physical measurements) can predict the embryos that are most likely to lead to successful implantation (**Fig. 4**). These data demonstrate that reproductive aging imposes a biomechanical checkpoint on embryonic morphogenesis and implantation (**Fig. 5**).

**Fig. 5.**
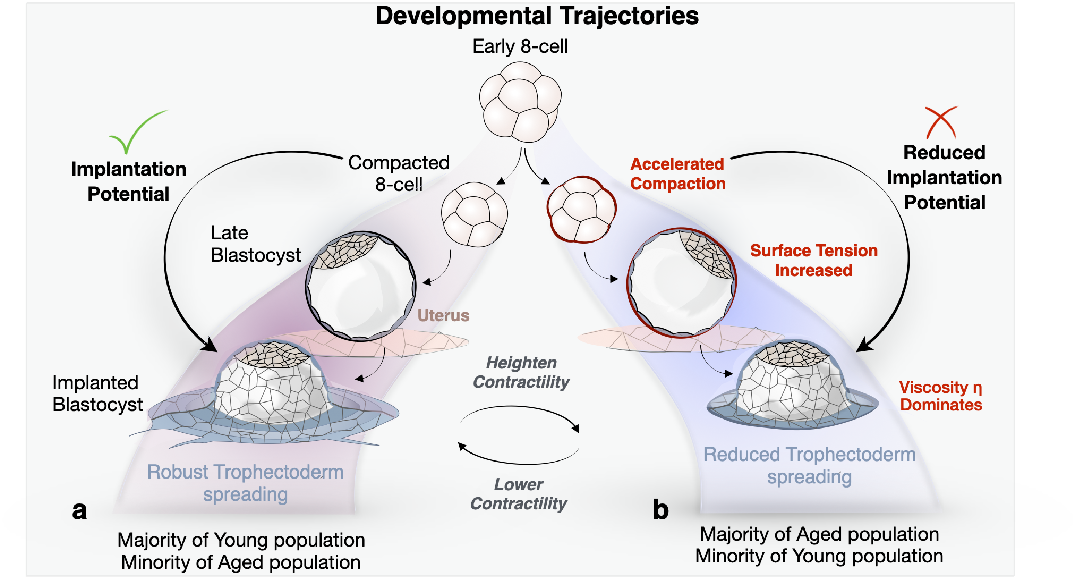
Summary Figure. Contractility shifts the proportion of implantation-competent embryos in the Young and Aged maternal condition. a) Schematic of developmental trajectory in the embryos that are competent to implant. Compaction proceeds normally at ∼ 9 h, followed by blastocyst formation and implantation to yield robust trophectoderm spreading. During wetting in these embryos, tissue-substrate adhesion forces dominate to produce robust trophectoderm spreading. The majority of Young embryos and minority of Aged embryos display this spreading behavior. Embryos that are incompetent to implant can be shifted to exhibit efficient wetting through lowering contractility. b) Schematic of developmental trajectory in the embryos that are incompetent to implant. Compaction is accelerated on average 2 hours faster due to aberrant contractility, that acts to increase surface tension at the blastocyst stage. Upon adhesion, tissue viscosity dominates to reduce wetting and slow trophectoderm spreading and implantation. The majority of Aged embryos and minority of Young embryos display this spreading deficiency. This spreading defect can be phenocopied in implantation-competent embryos by increasing contractile activity.

Understanding the cell biological origins of age-associated infertility is critical for improving embryo selection of the small pool of maternally aged embryos that are capable of uterine integration. Current embryo screening methods in IVF remain limited: morphological grading is subjective and often inconsistent between clinics, while preimplantation genetic testing for aneuploidy (PGT-A) does not provide a definitive diagnosis of the genetic status of the fetus proper^7,14,15,20,22,23,63^. Although chromosomal abnormalities are common with maternal age, even euploid embryos still frequently fail to implant, indicating that genetics alone cannot fully capture embryonic competence^7^. Mechanics may provide this missing dimension; cytoskeletal organization and tissue-level force balance not only regulate morphogenetic behaviors essential for implantation but are also increasingly recognized as upstream modulators of ploidy stability via cytoskeletal pathways^27,64^. Evidence from oocyte biology suggests that maternal age compromises cytoskeletal integrity, initial spindle positioning, and microtubule stability, potentially contributing to errors in chromosome segregation^8,9^. Thus, non-invasive mechanical screening of embryos has the potential to complement current morphology- and genetics-based assessments by capturing functional viability at the level of tissue behavior.

Our findings extend prior studies of mechanics-based morphogenesis defects for pre-implantation mouse and human embryos^62^. This previous work demonstrated that the 8-cell compaction stage represents a mechanical checkpoint where cortical tension and adhesion are integrated to establish blastomere flattening and robust blastocyst morphogenesis^57^. However, these obvious compaction defects are captured in existing embryo grading schemes^2^. Our work focuses on the critical process of implantation, for which there are no strong predictive metrics in the IVF field. The abbreviated compaction we observe in mouse embryos from aged females defines a distinctive mechanical signature that predicts implantation competence, forming the foundation for objective screening tools that may improve IVF outcomes for patients of all ages, given these results hold true in human embryos.

We identify hypercontractility as a core embryonic defect underlying age-related implantation failure, but the molecular basis of this mechanical abnormality is not known. The origins of this hypercontractility are likely multifactorial and may begin well before fertilization. While the aged uterus may be competent for embryo implantation, it could also serve as the source of alterations in egg quality. Importantly, the ovarian environment undergoes profound age-related remodeling^65,66^. The aging ovarian microenvironment is characterized by increased fibrosis, stromal rigidity, and altered extracellular matrix organization—changes that impair follicle development and oocyte quality^5^. Indeed, ovarian stiffness increases with age and directly influences follicle mechanics and oocyte maturation. These findings suggest that embryos from older females may inherit latent biomechanical liabilities established during oogenesis in a stiffened ovarian niche that could persist through cleavage divisions to compromise implantation. Alternatively, age-related changes in contractility may be a genetic hallmark of aging, as seen in other contexts such as stem cell niches^67^.

Beyond the potential for our work to identify the small subset of embryos from aged mothers that are competent for implantation, our studies open multiple translational opportunities for embryonic rejuvenation. Our pharmacological rescue experiments with Ripasudil suggest that mechanical defects may be reversible. This raises the possibility that therapeutic interventions — via targeted small molecules, culture conditions, or biomaterial cues that can “reset” contractility — could be implemented during the culture period in the IVF clinic to restore implantation competence in embryos from older mothers. Indeed, previous work showed that aged mouse oocytes, when transplanted to ovarian follicles from younger female mice, show markedly improved pregnancy success^68^. Additionally, reducing ovarian microenvironment stiffness via antifibrotic drugs is also associated with improved developmental outcomes^69^.Together, these data highlight the potential for rejuvenation in reproductive aging, offering renewed hope for rescuing developmental potential in embryos of older women. Our work potentiates similar translational tools that not only identify implantation-competent embryos but also build platforms for embryonic rejuvenation. Together our work, which is rooted in biophysics and mechanics, reveals a potential strategy to tackle age-related infertility to improve reproductive equity for older women.

## Materials and Methods

### Public Data on IVF Uterine Transfers

Surrogacy data was collated from the Human Fertilisation and Embryology Authority (HFEA)^24^. HFEA data was chosen for it being the most complete IVF dataset that included uterine transfers with aged female surrogates from aged female egg donors. Uterine transfers were selected and binned based on the age of the surrogate female and the age of the embryo donor. Young women were considered less than 35 years of age and aged women were considered over 35 years of age. Only eggs that were donated within specific binned maternal age categories were considered for downstream analyses. In this way, we excluded patient’s own eggs. Live birth rates from years 2012 to 2022 were then calculated as the birth rate per number of embryos transferred. Live birth rates were then averaged over the decade to give an averaged live birth rate for each transfer condition.

### Animal husbandry and Mouse lines

B6C3F1/J males were acquired from The Jackson Laboratory (Cat#100010) at 6 weeks of aged and utilized as male studs for breeding until 35 weeks of age. Vasectomized CD1 males were acquired from Charles River (Cat#022) at 6 weeks of age and 7-days post operation with sutures removed. CB6F1 females were used for breeding in two aged conditions: 6-8 weeks of age and 40 weeks of age. Young control CB6F1 females were acquired from The Jackson Laboratory (Cat#100009) and aged CB6F1 female mice were acquired through the National Institutes of Aging or acquired from the Jackson Laboratory (Cat#100009) and aged in house. Mice were housed on a 12 hour light:dark cycle and animal facilities were maintained at 21C. All animal work was carried out at the University of California, San Francisco (UCSF). Animal husbandry was performed in accordance with the UCSF Laboratory Animal Resource Center (LARC) guidelines. All experimental procedures were approved by the UCSF Institutional Animal Use and Care Committee (IACUC Protocol AN197455).

### Embryo Collections

Females were superovulated with 5IU PMS-G (Ilex A22721) at Day 1 and 5IU hCG (Ilex A225005) on Day 3, upon which they were placed with male studs for breeding. Copulation plugs were assessed on Day 4, and oviducts were surgically removed on Day 5 to collect embryos at E1.5 (2-cell stage). Oviducts were manually flushed using a blunt needle with EmbryoMax KSOM-AA Medium (EMD Millipore Cat#MR-101-D) to isolate embryos. Only phenotypically healthy 2-cell staged embryos were used for downstream assays; embryos were discarded if they showed severe fragmentation or malformation. Embryos were then washed in consecutive drops of KSOM-AA covered in Fijifilm light mineral oil (Fujifilm Cat#9305) before moving them to precalibrated drops of KSOM-AA under mineral oil. Embryos were incubated for culture in 37C, 5% CO_2_ conditions.

### NSET Procedures

Female surrogate mice of specified ages were superovulated as stated above. Embryos were collected at E1.5 and cultured until E3.5 Blastocyst stages. Recipient females were mated in parallel with vasectomized males and checked the next day for copulation plugs to confirm mating. Pseudopregnant females were timed for downstream use for NSET for 2.5 days post-coitum. E3.5 blastocysts were pooled and transferred using mNSET Devices for Mice (ParaTechs Corporation Cat#60010). Blastocysts were transferred in groups of 7-12 embryos to each recipient female to not overload each uterine horn. Surrogate females were monitored for distress for 3 days and again assessed for litter births 21 days post NSET procedure. NSET success was assessed as number of pups born divided by transferred blastocysts to calculate percentage of live births.

### In vitro Implantation Assays

In vitro implantation assays used were optimized from previously published reports^40^. In brief, embryos were collected at E1.5 and cultured until E3.5 blastocysts. Blastocysts were denuded using Acidic Tyrode’s solution (Sigma-Aldrich Cat# MR-004-D) until the zona pellucida was dissolved. Embryos were then transferred to KSOM-AA to deactivate any leftover Tyrode’s solution. In the event the zona pellucida was not fully removed, it was mechanically removed using a blunt glass pipette and manual pipetting. Blastocysts were then transferred to a 24-well plate (Corning, Cat#3526) with IVC media with 1:2000 dilution of SPY650-FastAct (Cytoskeleton Cat#CY-SC505) to begin incubation at D0. IVC media contained 1mg/mL Progesterone (Sigma-Aldrich Cat#P0130), 10μm Beta-estradiol (Sigma-Aldrich Cat#E2758-250MG), and 50mM N-Acetyl-L-cysteine (Sigma-Aldrich Cat#A7250) in advanced DMEM/F-12 (Life Technologies Cat#12634-010) supplemented with 25% ESC-grade Fetal Calf Serum (Life Technologies Cat#10439001), 1% ITS-X (Life Technologies Cat#51500-056), 1% Glutamax (Life Technologies Cat#35050-061), and 0.25% Pen/Strep (Gibco Cat#15140122). On Day 1, blastocysts were subsequently transferred to 6-well plates or Chamlide Chambers (Live Cell Instruments Cat#CM-M25-4) containing collagen-coated Polyacrylamide gels with fresh IVC media and SPY650-FastAct to enable live imaging of embryo implantation and/or Traction Force Microscopy. For immunostaining to analyze trophectoderm outgrowth, embryos were fixed as described in specific experimental workflows.

### Drug treatments

Embryos were treated with the cell-permeable drugs listed below starting at the late 2-to 4-cell stage before downstream applications. Control and drug-treated embryos were cultured each in a well from a 24-well plate (Corning, Cat#3526) in the absence of light mineral oil. Embryos were treated with equilibrated KSOM-AA and/or IVC media containing 1μg/mL RhoA Activator II (Cytoskeleton Cat #CN03) or 10μM Ripasudil (Selleck Chemical Cat#50-136-6407). Control embryos were treated with the same amount of vehicle (DMSO or water). Drugs were reconstituted according to manufacturer’s instructions before dilution to working conditions in KSOM-AA.

### Immunofluorescence

For blastocysts, embryos were placed in microbubbles with stated reagents on Greiner Bio-One Terasaki Plates (Cat#07-000-658) for all staining protocols. Gel-implanted embryos on glass coverslips were stained in 6-well dishes for all staining protocols. Embryos and coverslips were fixed in 4% PFA (Electron Microscopy Sciences Cat#15700) in PBS solution (Corning) at Room Temperature. Embryos were removed from PFA and then washed 3 times for 5 minutes each in PBS with 0.05% Tween-20 (PBST). Permeabilization was achieved through 0.3% Triton X-100 in PBS solution for 20 minutes and subsequently washed four times, 5-10 minutes each in PBST. Embryos were then blocked in blocking solution of 10% FBS diluted in PBST for 4 hours at room temperature. Antibodies were diluted in blocking solution and incubated overnight at 4C. Embryos were washed again three times for 10 minutes in PBST before being placed in secondary antibody solution with antibodies and DAPI diluted in blocking buffer. Embryos were incubated for 2 hours in secondary antibodies, washed four times in 10-minute PBST washes. Blastocysts were then transferred to a microbubble of PBS on a glass-bottomed MatTek dish (MatTek Corporation, P35G-1.5-20C) for downstream imaging. For gel-implanted embryos, glass coverslips were mounted with 10uL ProLong Gold (Invitrogen Cat# P36930) before being cured and sealed with nail polish. Immunostained blastocysts and coverslips were then stored at 4C before imaging.

### Antibodies

Primary antibodies used were: Rabbit pMLC (1:200; Cell Signaling Cat#3671S), Mouse ZO-1 clone 1A12 (1:200; Invitrogen Cat#33-9100), Goat ZO-1 (1:200; Invitrogen Cat#PA5-19090), Rabbit VD7 (2mg/ml, a kind gift from Dietmar Vestweber lab), Rabbit ppMLC (1:200; Cell Signaling Cat#3674T), Mouse Alpha-catenin clone 7A4 (1:500; Thermo Fisher Cat#13-9700), Rabbit NMIIA (1:200, BioLegend Cat#909802) Rat NMIIB (Abcam Cat# (EPR22564-23), Rabbit NMIIB (BioLegend Cat#909902). Secondary antibodies used were Alexa Fluor Goat anti-Mouse 647 (1:300, Invitrogen Cat#A21235), Alexa Fluor Goat anti-Mouse 568 (1:300, Invitrogen Cat#A11031), Alexa Fluor Goat anti-Rabbit 568 (1:300, Invitrogen Cat#A11055), Alexa Fluor Donkey anti-Rabbit 647 (1:300, Invitrogen Cat#A31573), Alexa Fluor Phalloidin 488 (1:100; ThermoFisher Cat#A12379), and DAPI (1:100, Sigma-Aldrich Cat#MBD0015).

### Calculating Junctional Intensity Levels of Tension-associated proteins

Junctional intensities of NMIIA, NMIIB, pMLC, alpha-catenin, and VD7 were calculated in Imaris and Excel. Surface renderings were performed using machine learning segmentation in the ZO-1 channel as a reference to isolate total junctions of the blastocyst. Junctions were virtually cut at cellular vertices to further isolate single cell-cell junctions along their length. Blastocysts’ junctions were then binned based on their presence at either polar or mural trophectoderm regions using the classification tool. Mean intensities of constituent channels were extracted using Imaris Statistics. Ratiometric values was then calculated for each junction using excel. For myosin levels, all channels were normalized to ZO-1 intensity levels. VD7 intensities were normalized to total alpha-catenin levels.

### Cell shape analysis

Mural Trophectoderm cell shapes were taken from fixed samples using the ZO-1 imaging channel. Imaging stacks were resliced using FIJI’s reslice tool to orient the bottom 1/3^rd^ region of the mTE and projected into a single en face plane. Image analysis to isolate cell shapes was performed in Python to extract cell perimeters and area only in the middle region of the 3D projection so as to not include any potentially stretched cells at the periphery.

### Calculating Trophectoderm Spreading

To analyze trophectoderm spreading, the 650-FastAct channel was used. Only the plane of tissue-substrate adhesion was used to calculate spread area. 650-FastAct was then thresholded in ImageJ to generate an actin mask and then particle analyzed to generate distinct Regions of Interest for each timepoint. ROIs were then measured in ImageJ and values were exported to Excel. The start of implantation was marked as T0 with the appearance of protrusive cells that made a stable blastocyst-substrate attachment that then expanded into an outgrowth. To calculate contact angles during Trophectoderm spreading, z-stacks were resliced in FIJI using the reslice tool, after which they were measured using the angle tool. Embryos were measured for multiple contact angles in 2 cross sections to generate an average contact angle for the embryo as it implants.

### Micropipette Aspiration Assay and Analysis

Micropipette Aspiration Package with −69mbar settings (Fluigent Cat#E-MASPI01) was mounted on a Leica DMi8 manual microscope with Integrated Modulation Contrast and Phase Contrast. Imaging was performed using a 40x HI Plan objective with 0.5 Numerical Aperture. Images were acquired using Leica Software at 0.2 millisecond intervals. Pressure was calibrated by using small 10nM spherical silica beads (a kind gift from Zev Gartner’s lab) that was mixed with a KSOM microbubble sitting immediately next to the measurement microbubble containing the embryos. Some particle beads were moved into the measurement bubble and zero pressure was calculated based on the point where there was no flow of the beads inwards or outwards of the micropipette. Using a glass micropipette with defined Radius R_p_, we non-invasively applied a pressure P_c_ to cells that exhibit a curvature of 1/R_c_. Micropipette glass (Biomedical Instruments) with diameters of 12μm and 14μm were used for compacting 8-cell stage embryos and E4.5 Blastocysts, respectively. Blastocysts were excluded from analysis if the experimental blastocyst deflated before measurements could be completed. Pressure values were recorded using Fluigent’s Oxygen software in parallel with live imaging recordings from the Leica software to match pressure values with cell aspiration. Using FIJI, we manually drew a circle around the target cell or trophectoderm region to analyze the radius of curvature. We used the linescan function to draw a line of the aspirated material into the micropipette to calculate Rp. Using the Young-Laplace equation, we quantified cellular surface tension γ_cm_ before and after the onset of compaction at the 8-cell stage. We then used Excel to calculate surface tension values after Pc was extracted from the Oxygen recordings.

### Calculating Compaction Metrics

For initial measurements of developmental tempo, embryos were imaged using either the spinning disk confocal or light sheet microscope. We then calculated developmental tempo as the dwell time within each cell cycle for each embryo. Namely, the timepoint between the resolution of cell division from the previous cell cycle and the initiation of the next subsequent cell division. Cell division was assessed based on the early bulging of the constituent cell and resolution of division or furrow ingression. Timepoints within the timelapse acquisition was marked in Excel and then calculated for dwell timing. To assess compaction metrics’ predictive power, embryos were indexed by plating individual embryos in a single square within 155μm Nylon Mesh Filters (Tisch Scientific Cat#ME17300) that were immobilized via Silicone sealant (Rutland Cat#76) on a Glass bottom 35mm Petri Dish with No. 1.5 coverglass (MatTek Cat#P35G1514). Developmental tempo was similarly assayed as above, in addition to measuring blastomere contact angles. At the start and endpoint of the cell cycle, angles between multiple blastomeres were measured in a single plane in Fiji using the angle tool. Multiple angles were then averaged to yield the average embryonic contact angle for the entire embryo to reflect its mechanical state.

### Live Imaging

For In Vitro Implantation Assays, E3.5 embryos were denuded with Acidic Tyrode’s solution and cultured in IVC media with 1:3000 dilution of 650-FastAct (Cytoskeleton Cat#CY-SC505). Upon imaging implantation, embryos were transferred to fresh IVC including fresh 650-FastAct dilution for subsequent imaging. For imaging preimplantation morphokinetics, embryos were collected at E1.5 and cultured in KSOM-AA and 1:3000 dilution 650-FastAct for direct downstream imaging. Imaging was begun after 4 hours to enable uptake of the 650-FastAct into cytoskeletal structures. Alternatively, embryos were labeled using microinjection technique. mRNA from the vector pRN3P_membrane_EGFP (Addgene Cat#139402) was isolated using mMESSAGE mMACHINE T7 Transcription Kit (ThermoFisher Cat#AM1344) according to manufacturers specifications. Membrane_EGFP RNA was injected at E1.5 into each blastomere of the embryo at concentrations of 100ng/uL. Embryos were placed in an equilibrated and heated droplet of KSOM-AA on a depression slide covered by light mineral oil. Injections were performed using the Eppendorf Femtojet microinjector mounted on Leica DMi8 manual microscope with Integrated Modulation Contrast using a 40x HI Plan objective with 0.5 Numerical Aperture.

### Traction Force Microscopy

Traction force microscopy was performed as described previously^49,50^. Briefly, either 25mm round glass coverslips (Electron Microscopy Sciences Cat#72223-01) or 18mm round glass coverslips (Fisher Scientific, Cat#12541005) were cleaned with plasma for 2 minutes using a Plasma Cleaner (Harrick Plasma, Cat#PDC-001). Glass coverslips were then bathed in 2% 3-aminopropyltrimethoxysilane (Sigma Aldrich cat#281778) in isopropanol for 10 minutes, washed 4x for 10 minutes in ddH2O, and then dried overnight covered in tinfoil in a dust-free environment. Coverslips were then treated with 1% glutaraldehyde in ddH2O for 30 minutes, washed 3x for 10 minutes in ddH2O, and again dried overnight in tinfoil a dust-free environment. PAA gel was prepared using a 16kPa Shear modulus Standard Solution with a final working solution of 12% Acrylamide (BioRad Cat#161-0140) and 0.15% Bis-Acrylamide (Fisher Scientific Cat#BP1404-250) and polymerized using 10% Ammonium Persulfate (Fisher Scientific Cat#BP179-25) and TEMED (Fisher Scientific Cat#BPP150-20). Traction Force Microscopy gels additionally contained 200nm fluorescence microspheres (Invitrogen Cat#F8807 or Cat#F8811). Collagen ECM-coupling to the PAA gel was achieved by UV crosslinking of Sulfo-SANPAH (Fisher Scientific Cat#PI22589) diluted in DMSO and incubation with 2mg/mL Collagen (Corning Cat#354236) overnight in 4°C. Gels were rinsed 5-10x in sterile PBS and UV irradiated in a germicidal lamp for 10 minutes before embryo plating in equilibrated IVC media. Embryos were plated onto the gels at E4.5 and imaged prior to implantation attachment to establish baseline reference images of the fluorescent beads. Embryos were pooled by maternal condition and placed in separate wells of the Chamlide, or when both conditions were pooled together, Young embryos were injected at the 2-cell stage with membrane-eGFP and identified post-imaging to denote maternal condition. Traction forces were analyzed using code written in Python (https://github.com/OakesLab/TFM) according to previously described work^70^. Prior to image processing, images were flat-field corrected, with the reference bead image aligned to embryo-attached bead images. Bead displacement was calculated using an optical flow algorithm in OpenCV (Open Source Computer Vision Library, https://github/itseez/opencv) with a window size of 8 pixels. Traction stresses were calculated using the Fourier Transform Traction Cytometry approach^71^ with a regularization parameter of 3.35 *×* 10^−6^. The total contractile work, or strain energy, was calculated as one half the dot product of the traction stress and displacement vectors, summed over the area of the trophectoderm.

### Fit to experimental data

We fit the model solutions (see Supplementary Note) to the experimental data on the evolution of the projected area of the blastocysts, *A*(*t*) (Supp. Fig. 2e). By numerically solving Eq. (13) and Eq. (14) in the Supplementary note, we obtain the radius evolution *R*(*t*), from which we obtain *A*(*t*) = *πR*^2^. We fit this quantity to the experimental data, with the initial contact angle *θ*_0_, the traction magnitude *T*_0_, and the tissue viscosity *η* as fitting parameters. We estimate the remaining parameters directly from our experimental results. First, we obtain the initial radius *R*_0_ directly from the area measurements as 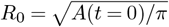. Second, we use the values of the tissue surface tension measured by micropipette aspiration (Fig. 3c), which are *γ*_young_ = 1.4 *±* 0.1 mN/m and *γ*_aged_ = 2.1 *±* 0.1 mN/m. Third, we estimate the traction decay length *L*_c_ as follows: The large fluctuations of the experimental traction fields make it difficult to reliably fit the measurements with the model. Consequently, to estimate *L*_c_, we locate the minimum (most negative) radial traction near the blastocyst boundary, *T*_min_, and we obtain *L*_c_ as the distance inward at which traction decreases to *T*_min_*/e* (Supp. Fig. 2c). Using this method, we obtain the values shown in Supp. Fig. 2c for embryos from young and aged mothers. Based on these results, we set *L*_c_ = 15 *μ*m. This choice reflects a compromise: Rather than discarding the higher values in the long tail of the distribution, we select the upper quartile as a more representative estimate than the median. With these parameter estimates, we obtain fits such as shown in the examples (Supp. Fig. 2e). From them, we obtain values of the initial contact angle *θ*_0_, which agree well with the experimentally measured contact angles at the moment of attachment (T0 in Supp. Figs. 2d, 2e). We also obtain the average values of the active traction magnitude *T*_0_ and the tissue viscosity *η* shown in Fig. 3g,h. From the averages, we excluded outliers using an interquartile range (IQR) criterion.

### Microscopy

A spinning disk confocal was equipped with a Nikon Ti2-E body and configured with a CrestOptics X-Light V3 confocal spinning disk system, a Lumencor Celesta light engine, Nikon 10x CFI Plan Apo Lambda objective, an Okobox temperature and CO_2_ controlled environment, and a Photometrics Kinetix sCMOS camera. Light sheet imaging utilized a Luxendo TruLive3D microscope equipped with a Nikon 25x 1.1NA detection objective and Hamamatsu Flash 4 cameras with a final magnification of 31.25x. Excitation was performed with Omicon and OBIS lasers with an effective light sheet FWHM of 1.6um thick. All images were acquired using LuxBundle software version 4.3.4 and analyzed in FIJI.

### Statistical Analysis

Statistical analysis was performed in Excel, GraphPad Prism, and Matlab, to establish statistical significance under the defined specific experimental conditions. Outliers were identified using the ROUT method (Robust regression and outlier removal) with a false discovery rate of Q=5% and excluded from subsequent analysis. The statistical test used, exact n values, and definition of replicates are provided in the figure legends. N represents the experimental replicates, and n represents the number of embryos used in the indicated maternal or drug conditions. Data were analyzed for significance with ****=p<0.0001, ***=p<0.001, **=p<0.01, and *=p<0.05, which is indicated in the figure and its respective legend, as determined by respective tests described in the figure legends.

## Supporting information

SMov1_YvsA-spreading

SMov2_E4.5-MPA

SMov3_YvsA-TFM

SMov4_YvsA-Compaction

SMov5_E2.5-MPA

SMov6_IndexedCompaction

## Author Contributions

KEC, ODW, and DJL conceived of the study. KEC collected and analyzed data. MJFO and RA developed the theory and fitted it to the data. PWO assisted with optimization and analysis of TFM experiments. KEC, ODW, MJFO, and RA wrote the original draft. All authors edited the manuscript. All authors approved the final version of the article.

## ACKNOWLEDGEMENTS

We thank the entire Weiner Lab for helpful suggestions insightful comments, and critical reading of the text. We thank Sneha Rao and Elysse Phillips of the Weiner Lab with critical help setting up the mouse embryo as the model system of use, in addition to setting up experimental frameworks. We thank Dane Maxfield (Bruker) for generous use of the TruLive3D Lightsheet Microscope. We thank UCSF’s Center for Advanced Light Microscopy (CALM) for use of its core microscopes. KEC was supported by Jane Coffin Childs Postdoctoral Fellowship, Burroughs Wellcome Fund Postdoctoral Enrichment Program, Bakar Aging Research Institute Postdoctoral Fellowship, and Burroughs Wellcome Fund Career Award at the Scientific Interface. ODW was supported by GM118167, a Sandler Program for Breakthrough Biomedical Research Grant, and a collaborative grant with Altos. PO was supported by GM148644. DJL was supported by 1R01GM122902, 1R01ES023297, and the Global Consortium for Reproductive Health through the Bia-Echo Foundation GCRLE-0123. MJFO and RA acknowledge funding from the Max Planck Society. RA acknowledges funding from the European Union’s ERC Starting Grant “Living Fluctuations” No. 101114584.

## Supplemental Figures

**Supplemental Figure 1.**
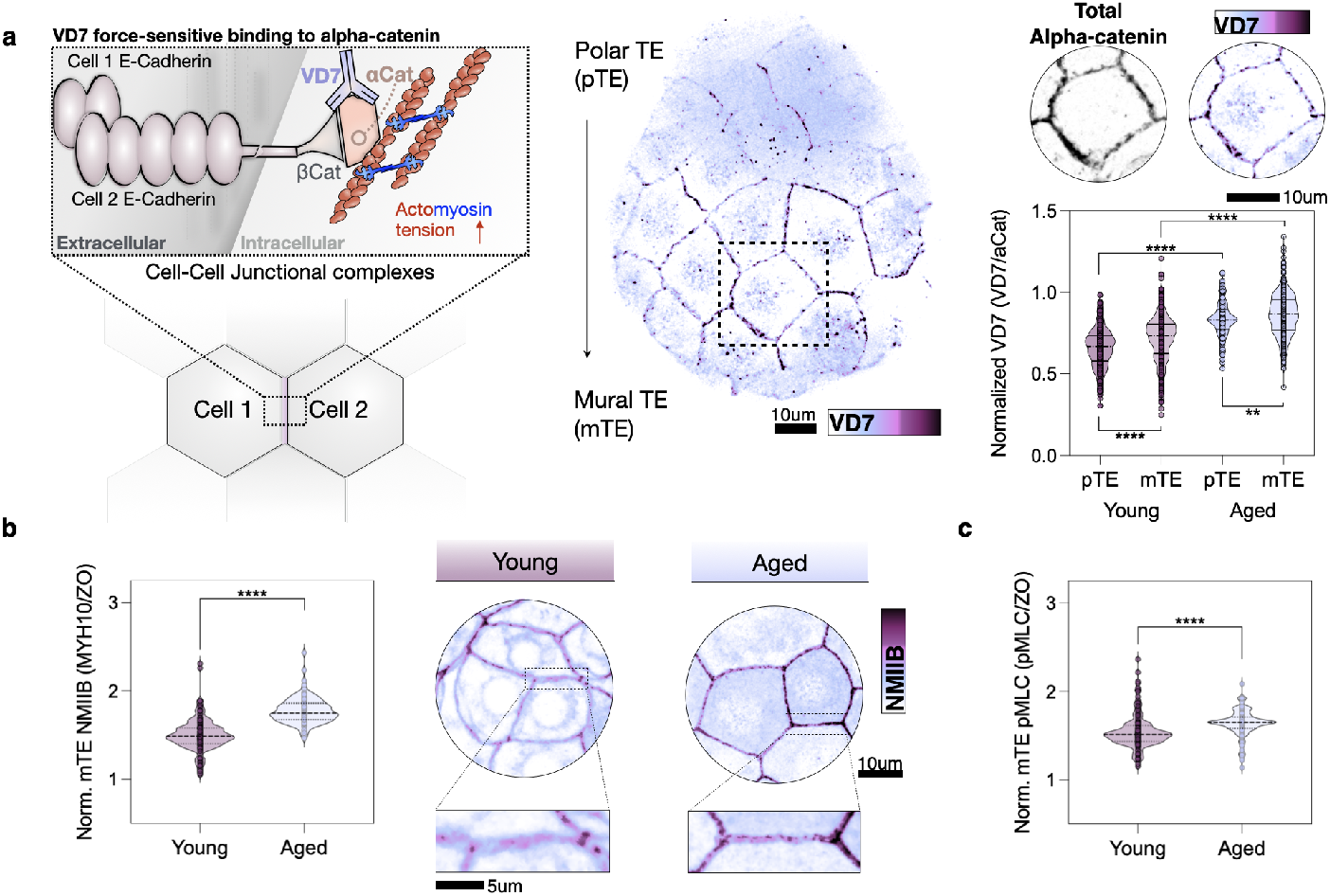
Maternally aged embryos show increased mechanochemical signaling. a) Immunostaining of late E4.5 blastocysts with the tension sensor, VD7. The VD7 antibody is an engineered antibody sequence based on vinculin’s binding association to alpha-catenin. Contractility at cell-cell junctions creates a force-dependent conformational change in alpha-catenin proteins that exposes this epitope pocket to which vinculin binds. VD7 association is observed when E-cadherin-mediated adhesions experience tension, making it an indicator of tissue under tensile strain. We measured global junctional VD7 and normalized its levels to total alpha catenin to quantify relative junctional VD7 signal intensity. Blastocyst fixed and stained for tension sensor VD7, showing increasing junctional VD7 towards the mural trophectoderm region. Normalized VD7 (purple) was calculated over total alpha-catenin (black), showing that maternally aged embryos show higher junctional tensions both at the mural and polar trophectoderm regions. (Young N = 4, n = 386 pTE junctions, 532 mTE junctions, 10 blastocysts; Aged N = 3, n = 218 pTE junctions, 383 mTE junctions, 8 blastocysts). p values from Mann-Whitney test. Scale bar is 10μm. b) Plot showing Normalized junctional NMIIB staining in the mTE region, with higher junctional accumulation in the aged condition (left) (Young N = 2, n = 1,610 mTE junctions, 17 blastocysts; Aged N = 2, n = 935 mTE junctions, 14 blastocysts). p values from Mann-Whitney test. Scale bar is 10μm. Representative images of NMIIB staining (purple) in both maternal conditions. Scale bar is 5μm and 10μm. c) Plot showing Normalized junction pMLC staining in the mTE region, with higher junctional accumulation in the aged condition (Young N = 2, n = 1,610 mTE junctions, 17 blastocysts; Aged N = 2, n = 935 mTE junctions, 14 blastocysts). p values from Mann-Whitney test.

**Supplemental Figure 2.**
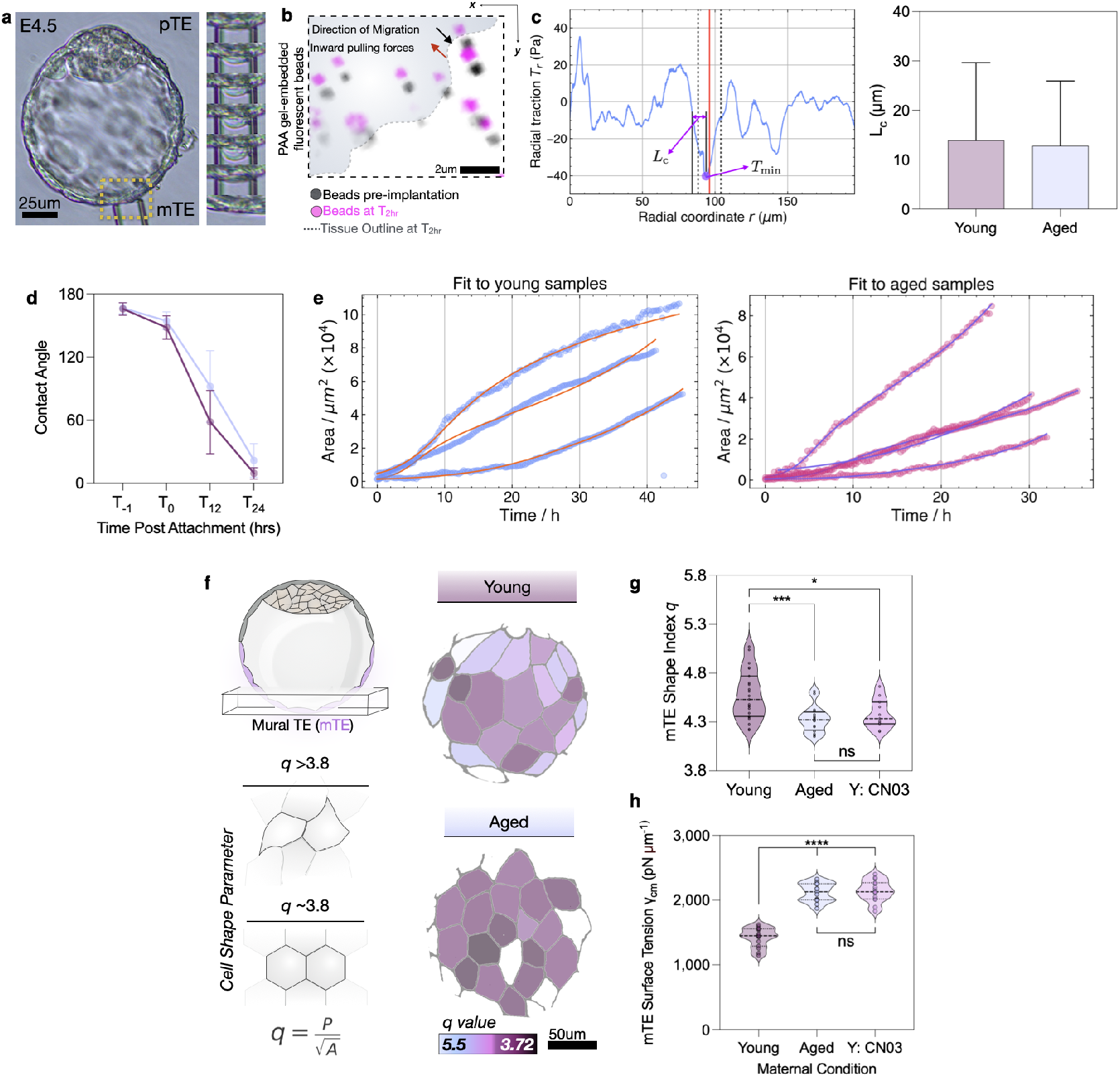
Aged embryos show increased viscosity in the wetting model. a) Representative image of an E4.5 aged blastocyst undergoing Micropipette aspiration assay at the mural trophectoderm region (left) with zoomed-in timelapse of the yellow highlighted region showing the progression of tissue aspiration over time (right). Scale bar is 25μm. b) Representative outline of a migrating trophectoderm front (shaded blue) in the direction of the black arrow, with pulling forces going in the opposite direction. Black beads represent the reference image of the beads before implantation, with the pink beads representing the beads after 2 hours of implantation, together showing bead displacement as a function of tissue migration. Scale bar is 2μm. c) Example frame of a radial traction profile used to estimate Lc. The red line marks the blastocyst boundary, while the dashed lines indicate the window around the boundary used to account for uncertainties in image analysis. Within this window, the minimum traction is identified, and Lc is determined as the distance at which the traction decays to T0/e (left). Measurements of the length scale Lc extracted from the traction stress profiles (Young n = 589 profiles; Aged n = 642 profiles). Box plots show mean and SD (right). d) Measurement of blastocyst contact angles over 24 hours of implantation (Young N = 3, n = 41 blastocysts; Aged N = 3, n = 41 blastocysts). Dots represent mean with SD. e) Example of model fits to the blastocyst area expansion for both maternal conditions. Colored dots represent the area measurements. Solid lines indicate the model fits. f) Schematic of mTE analysis of q values (left) and representative images (right) of blastocyst cell segmentation as a function of cell shape parameter q in both maternal conditions. Aged conditions show more hexagonal cell shapes. g) Plot showing individual q values for cells in the mTE region of blastocysts from Young, Aged, and Young-CN03 treated embryos (Young N = 3, n = 384 cells, 27 Blastocysts; Aged N = 3, n = 239 cells, 19 Blastocysts; Young-CN03 N = 3, n = 221 cells, 22 Blastocysts). p values from Mann-Whitney test. h) Plot showing surface tension measurements of E4.5 blastocysts from Young, Aged, and Young-CN03 treated embryos (Young N = 3, n = 15 blastocysts; Aged N = 3, n = 15 blastocysts; Young-CN03 N = 3, n = 15 blastocysts). p values from Mann-Whitney test.

**Supplemental Figure 3.**
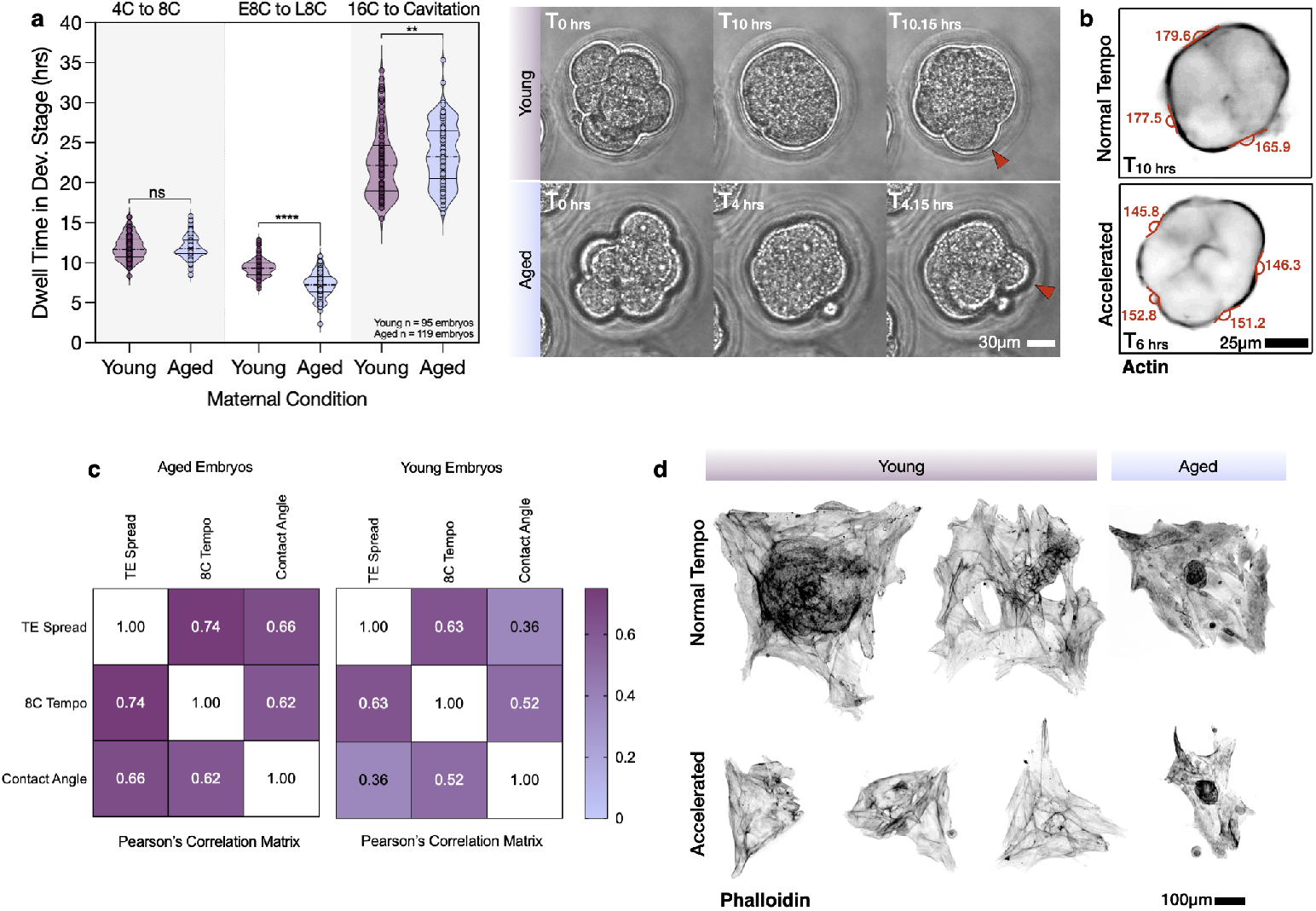
Compaction metrics determine implantation potential. a) Quantification of developmental tempo for all examined stages. Compaction undergoes an age-related abbreviation, while cavitation timing is lengthened (left) (N=4, Young n = 95 embryos; Aged n = 119 embryos). p values from Mann-Whitney test. Timelapse video (right) showing Young (top) and Aged (bottom) embryos undergoing compaction using Brightfield imaging. T0 marks the beginning of the 8-cell stage. Aged embryo undergoes a significantly accelerated compaction from the early to late 8-cell stage. Red arrows denote cell divisions, denoting the end of the 8-cell stage. Scale bar is 30μm. b) Representative images from a timelapse of embryos stained for 650-FastAct showing Normal and Accelerated compaction conditions. Red labels denote the calculated inter blastomere contact angles. Accelerated compacted embryos showed more acute angles than embryos with Normal tempo. Scale bar is 25μm. c) Pearson’s correlation matrix showing correlation between final trophectoderm spreading, 8-cell compaction tempo, and final compaction inter-blastomere contact angles for both maternal conditions. Both 8-cell tempo and contact angles show significant correlations to final trophectoderm spreading values (Young n=56 embryos; Aged n = 14 embryos). d) Representative images of implanted embryos stained for Actin that underwent Normal (top) and Accelerated (bottom) compaction rates. Aged embryos are denoted with light purple color. Embryos selected for normal compaction tempo exhibited robust trophectoderm spreading events in both maternal conditions. Embryos selected for accelerated compaction tempos showed reduced trophectoderm spreading in both conditions. Scale bar is 100μm.

## Supplemental Movie Legends

**Movie 1: Maternally aged embryos show reduced trophectoderm spreading**. Representative timelapse of Young (left) and Aged (right) implanting blastocysts over the 24 hours post-implantation. Blastocysts are stained for 650-FastAct and displayed as z-projection. Scale bar is 50 μm.

**Movie 2: Maternally aged embryos show increases in blastocyst surface tension**. Representative timelapse of Young (left) and Aged (right) E4.5 blastocyst undergoing Micropipette Aspiration assays. Scale bar is 50 μm.

**Movie 3: Traction force microscopy of aged embryos shows increased tissue-substrate contractility**. Actin stained for 650-FastAct (top) in Young (left) and Aged (right) maternal conditions showing the attachment plane of the mural trophectoderm. Calculated contractile traction stresses (bottom) show increased tissue-substrate contractility in the aged condition. Scale bar is 50 μm.

**Movie 4: Embryos in the aged condition undergo precocious compaction**. Representative timelapse of Young (left) and Aged (right) embryos transitioning from the 4-cell stage to the blastocyst stage. Aged embryos accelerate through the 8-cell stage and undergo precocious compaction. Scale bar is 50 μm.

**Movie 5: Micropipette aspiration of 8-cell stage embryos**. Representative timelapse of Early (left) and Late (right) 8-cell stage embryos undergoing Micropipette Aspiration Assay. Scale bar is 10 μm.

**Movie 6: Indexed embryos identify fast and slow compaction tempos**. Representative timelapse of indexed embryos on silicone grid undergoing development from the 2-cell to the 16-cell stage. Embryos are live stained for 650-FastAct. Zoomed in timelapses represent embryos with Normal and Accelerated developmental tempo. Scale bar is 100 μm.

## Supplementary Note 1: Active Wetting Model

### A. Active wetting model

Here, we provide a detailed description of our physical model of tissue spreading. Building on the approaches of Refs.^41,48^, our model explains tissue spreading as the wetting process of an active droplet. The droplet takes the shape of a spherical cap, with fixed 𝒱 and varying contact radius *R*(*t*) and contact angle *θ*(*t*) (Fig. 3b). For a droplet with a contact angle *θ <* 90^°^, spreading is driven by active traction forces **T**_a_ exerted by cells on the substrate, and it is opposed by the tissue surface tension *γ* (Fig. 3b).

In our model, active tractions are generated by cells that are in contact with the substrate and exhibit front-back polarity, which is described through the polarity field **p**(**r**, *t*). Following previous work^41^, we assume that the basal cell layer is unpolarized in the bulk and polarized at the edges, where we impose *p*_*r*_(*R, t*) = 1. The polarity field is then taken to follow relaxational dynamics, independent of the flows in the cell layer:

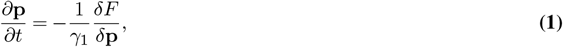

where *F* is the free energy of the polarity field. This free energy favors the unpolarized state **p** = 0 with a restoring coefficient *a*, and penalizes spatial variations of the polarity with a Frank constant *K*^72^:

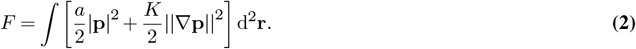

We assume that the polarity field relaxes much faster than the time scale of tissue flows, so that it is effectively at equilibrium. Thus, setting *∂***p***/∂t* = 0, the polarity field obeys

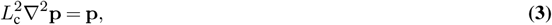

where 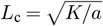 is the characteristic length over which the polarity field decays from its maximum at the tissue edge to zero in the bulk.

On the other hand, since tissue flows occur at very low Reynolds numbers, inertial forces are negligible. Consequently, the basal cell layer satisfies force balance among all relevant forces, which can be expressed as^41^

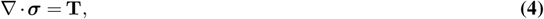

where *σ*(**r**, *t*) is the monolayer stress tensor and **T**(**r**, *t*) is the substrate traction. We model the monolayer stresses as that of a viscous fluid:

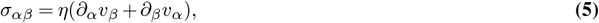

where *η* is the viscosity, which arises from cell-cell friction, and the Greek indices *α* and *β* denote the spatial components. Respectively, the substrate traction has contributions from friction, with a coefficient *ξ* and proportional to the monolayer velocity field **v**(**r**, *t*), and from active cell-generated forces, with magnitude *T*_0_ and proportional to the polarity field **p**(**r**, *t*):

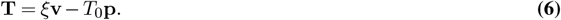

This model features two sources of dissipation: tissue viscosity and cell-substrate friction, which define the hydrodynamic screening length, 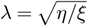. This length scale indicates how far stresses can propagate in the tissue before being damped by friction with the substrate. When *λ > R*, viscosity dominates, and substrate friction has only a weak effect on the velocity and stress fields. For epithelial monolayers, typical values of the screening length are *λ* ∼ 0.2 − 0.6 mm^41^, while the radii of monolayers in our experiments with mouse embryos rarely exceed 0.2 mm. Therefore, we assume that viscosity dominates over friction and neglect cell-substrate friction in the following analysis. With this assumption, the final expression for the force balance Eq. (4), after introducing the constitutive equation Eq. (5), is

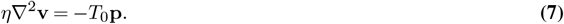

Finally, we specify the boundary condition for the tissue flow field **v**(**r**, *t*). Following Ref.^48^, and based on the physics of wetting, we use a generalized Young-Dupré condition to account for the role of surface tension at the tissue edge. This boundary condition balances the horizontal component of the surface tension *γ* with the internal stresses at the edge of the basal cell layer:

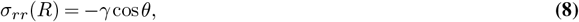

where *θ* is the contact angle.

### B. Traction and velocity profiles

Here, we solve the model for a circular basal cell layer of radius *R*, corresponding to the bottom surface of the spherical cap droplet. We first solve Eq. (3) to obtain the polarity field, and then use it in Eq. (7) to solve for the flow field.

#### B.1. Polarity field

We assume axial symmetry, so that the polarity field is radial: 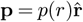. In polar coordinates, Eq. (3) then becomes

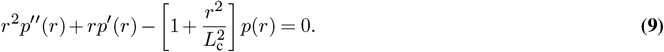

Imposing the boundary condition *p*(*R*) = 1, together with the symmetry condition *p*(0) = 0, the radial polarity profile

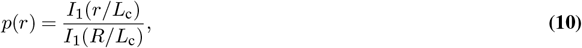

iswhere *I*_1_ is the modified Bessel function of the first kind and first order. Using this expression, and neglecting friction in Eq. (6) as discussed above, the traction profile follows directly as 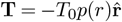.

#### B.2. Velocity field

Similarly, we assume the velocity field to be purely radial, 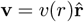. In polar coordinates, Eq. (7) then reads

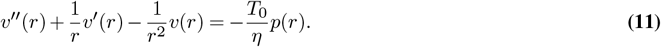

We solve this equation with the boundary condition Eq. (8), which can be written as *v*^*t*^(*R*) = −*γ/η* cos *θ*. Using also the symmetry condition *v*(0) = 0, we obtain

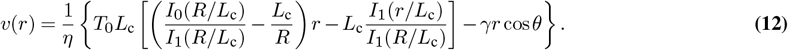

#### B.3. Spreading dynamics

Given the velocity field above, the monolayer radius evolves as 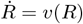. Since we model the blastocyst as a spherical cap (Fig. 3b) with a fixed volume 𝒱 = *πR*^3^*/*3(2 − 3 cos *θ* + cos^3^ *θ*)*/* sin^3^ *θ*, the contact angle *θ* must decrease as the tissue spreads. To simplify the expressions, we write 𝒱 = *πR*^3^*/*3*f* (*θ*), where we introduced the trigonometric function *f* (*θ*) = (2 − 3 cos *θ* + cos^3^ *θ*)*/* sin^3^ *θ*. Imposing that 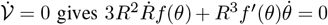, which relates 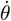 to 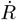. Thus, we finally obtain a system of two coupled differential equations that describe the spreading dynamics:

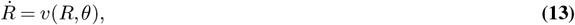

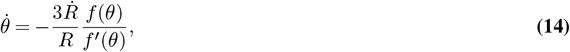

We solve these equations numerically to fit the experimental data on the evolution of the projected area of the blastocysts, as explained in the Methods section.

